# Stable and widespread structural heteroplasmy in chloroplast genomes revealed by a new long-read quantification method

**DOI:** 10.1101/692798

**Authors:** Weiwen Wang, Robert Lanfear

**Affiliations:** Research School of Biology, Australian National University, Canberra, ACT, Australia, 2601

**Keywords:** Single copy inversion, flip-flop recombination, chloroplast genome structural heteroplasmy

## Abstract

The chloroplast genome usually has a quadripartite structure consisting of a large single copy region and a small single copy region separated by two long inverted repeats. It has been known for some time that a single cell may contain at least two structural haplotypes of this structure, which differ in the relative orientation of the single copy regions. However, the methods required to detect and measure the abundance of the structural haplotypes are labour-intensive, and this phenomenon remains understudied. Here we develop a new method, Cp-hap, to detect all possible structural haplotypes of chloroplast genomes of quadripartite structure using long-read sequencing data. We use this method to conduct a systematic analysis and quantification of chloroplast structural haplotypes in 61 land plant species across 19 orders of Angiosperms, Gymnosperms and Pteridophytes. Our results show that there are two chloroplast structural haplotypes which occur with equal frequency in most land plant individuals. Nevertheless, species whose chloroplast genomes lack inverted repeats or have short inverted repeats have just a single structural haplotype. We also show that the relative abundance of the two structural haplotypes remains constant across multiple samples from a single individual plant, suggesting that the process which maintains equal frequency of the two haplotypes operates rapidly, consistent with the hypothesis that flip-flop recombination mediates chloroplast structural heteroplasmy. Our results suggest that previous claims of differences in chloroplast genome structure between species may need to be revisited.

**Significance Statement:** Chloroplast genome consists of a large single copy region, a small single copy region, and two inverted repeats. Some decades ago, a discovery showed that there are two types chloroplast genome in some plants, which differ the way that the four regions are put together. However, this phenomenon has been shown in just a small number of species, and many open questions remain. Here, we develop a fast method to measure the chloroplast genome structures, based on long-reads. We show that almost all plants we analysed contain two possible genome structures, while a few plants contain only one structure. Our findings hint at the causes of the phenomenon, and provide a convenient new method with which to make rapid progress.

## Introduction

Chloroplasts are organelles which are vital for photosynthesis. Most land plant chloroplast genomes are 120 – 160 kb in size (1, 2), and have a quadripartite structure consisting of a pair of identical rRNA-containing inverted repeats (hereafter referred to as IR) of ∼10-30kb divided by a large single copy (LSC) region of ∼80-90 kb and a small single copy (SSC) region of ∼10-20 kb.

Surprisingly, chloroplast genomes can exist two structural haplotypes differing in the orientation of single copy regions (3). The presence of this structural heteroplasmy has been confirmed in some land plants (3-6) and algae (7, 8), but its cause remains unknown. One hypothesis to explain the presence of structural heteroplasmy is known as flip-flop recombination (4). This hypothesis suggests that the large IRs could mediate frequent intramolecular recombination, resulting in the maintenance of roughly equal amounts of the two haplotypes differing only in the orientation of their single copy regions.

A better understanding of chloroplast genome structural heteroplasmy is important for a number of reasons. First, some recent papers have suggested that the orientation of the single copy regions differ between species (9-13), but Emery *et al* (14) pointed out that these studies seem to have overlooked the possibility that both orientations may coexist in a single individual. Second, the relationship between the structure of the chloroplast genome and the existence or otherwise of structural heteroplasmy remains poorly understood. For example, if flip-flop recombination causes chloroplast structural heteroplasmy, then it seems likely that the presence of two long IRs may be a pre-requisite for the existence of heteroplasmy. Consistent with this, no heteroplasmy was observed in a chloroplast genome with highly reduced IRs (15), but the generality of this observation remains to be tested. Third, if heteroplasmy is the norm rather than the exception, then this may represent a challenge for the assembly of chloroplast genomes, which may be easily overcome by simply allowing for the existence of two structural haplotypes during the assembly process. A large-scale analysis across many different species has the potential to provide a more complete picture of chloroplast heteroplasmy, further elucidating this fascinating phenomenon and potentially improving genome assembly and inference. In this study, we perform this large-scale analysis using a new method which we developed to quickly and conveniently quantify structural heteroplasmy in chloroplast genomes from long-read sequencing data.

Currently there are two methods to detect different structural heteroplasmy in chloroplast genomes: Bacterial Artificial Chromosomes (BAC)-End-Sequence (BES) (5), and restriction digests (4). For the BES method, one first constructs BAC libraries in which large pieces of chloroplast DNA are inserted into bacterial chromosomes. One then uses the BAC libraries to sequence short fragments of both ends of many such pieces of chloroplast DNA to ascertain which chloroplast structures exist in a given sample. BES reads can only provide information on chloroplast genome structure when a single read covers an entire IR region, with one end in LSC region and another end in the SSC region. Although a very useful method, BES reads may be limited for detecting highly atypical chloroplast genome structures, because only a short fragment from each end of a read is actually sequenced. For the restriction digestion method, one uses restriction enzymes to digest the chloroplast genome, and then decodes the chloroplast genome structure by studying the distribution of the resulting fragment lengths using agarose gel electrophoresis. Using this approach, Stein *et al* (4) found that the IR region was present in four different fragments which could be separated into two groups for which the sum of lengths was equal. They therefore concluded that chloroplast genome contained two equimolar isomers, which differed in their single copy orientation. The restriction digest method has provided much of the existing information on chloroplast genome heteroplasmy, but it is limited by the availability of suitable restriction sites (which may be unknown in some species), and requires a labour-intensive hybridisation step to infer chloroplast genome heteroplasmy. Other methods have also been proposed, such as the use of PCR to attempt to amplify diagnostic regions of the chloroplast genome. However, it is not easy to generate PCR fragments which are longer than the IR region (usually 10-30 kb), so while PCR-based methods could only work well with short IRs (e.g. of a few hundred bp) (15), their general application is likely to remain limited. Furthermore, PCR-mediated recombination could result in false positives when using PCR-based methods (16, 17). In short, current methods for detecting structural heteroplasmy in chloroplast genomes are labour- and time-intensive, which makes them difficult to employ for broad-scale studies. Current methods have been applied in a relatively small number of land plant species (*Beta vulgaris* (6), *Musa acuminata* (5), *Phaseolus vulgaris* (3), *Osmunda cIaytoniana, Osmunda cinnamomea* and *Osmunda regalis* (4), and some species in the Pinaceae (15)), and as a result our understanding of the patterns and causes of structural heteroplasmy in chloroplast genomes remains somewhat limited.

In this study, we develop and apply a fast, simple, and cheap method to detect and quantify structural heteroplasmy in chloroplast genomes using long-read sequencing. This method requires just a single DNA extraction and sequencing run, followed by a simple bioinformatic analysis for which we provide free and open-source code (https://github.com/asdcid/Cp-hap). Our approach, which we call Cp-hap, relies on sequencing individual DNA molecules longer than the length of the IR region (roughly 10-30 kb in most species). Sequencing reads of this length are commonly obtained in great abundance when sequencing whole-plant DNA extractions on the Oxford Nanopore MinION device (18, 19). Our method sifts through these long sequencing reads to find those that uniquely match one of the 32 uniquely-identifiable structural haplotypes (Fig. S1), and then uses the abundance of these reads to estimate the relative abundance of each haplotype (details see Materials and Methods). Since many new plant samples are being sequenced with long-read sequencing technology, this method has the potential to provide many insights in the coming years, simply as a by-product of existing sequencing efforts.

We use Cp-hap to quantify the abundance of chloroplast genome structural haplotypes in 61 plant species. Our results suggest that most land plants contain the two commonly-found chloroplast genome haplotypes in equal abundance whenever the chloroplast genomes contain a pair of large IRs. However, when the IR is small (e.g. only a few hundred bp in size) or repeat regions are not inverted but positioned, we find only a single haplotype. Finally, we show that multiple samples taken from a single individual all contain the same two haplotypes in equal abundance, suggesting that the process which maintains equimolar concentrations of the two haplotypes operates relatively rapidly, and consistent with the suggestion that flip-flop recombination is the underlying cause.

## Results

### Data collection

We searched the National Center for Biotechnology Information (NCBI) Sequence Read Archive (SRA) database, and selected 90 land plants species which met our length and data amount requirement (see Materials and Methods) to analysis (Dataset S1). Two of these species (*Herrania umbratica* and *Siraitia grosvenorii*) lacked suitable chloroplast reference genomes, so we therefore assembled these two chloroplast genomes *de-novo* (see Materials and Methods). The chloroplast genomes of *Herrania umbratica* and *Siraitia grosvenorii* were 158,475 bp and 158,757 bp in size, consisting of a LSC (88,369 bp and 87,625 bp), a SSC (19,040 bp and 18,556 bp), and two IRs (25,533 bp and 26,288 bp), and encoding 134 and 133 genes respectively (Fig. S2).

### Chloroplast structural haplotypes across the plant tree of life

For these 90 species, we obtained adequate long-read sequencing data to assess chloroplast genome structural heteroplasmy for 61 land plants species. Across all 61 species, we find just three structural haplotypes (Fig. 1, Table 1). All 58 of the Angiosperms in our dataset are heteroplasmic, and contain roughly equal frequencies of haplotypes A and B (binomial test for departure from equal frequency, p value >0.05 in all cases). The remaining three species are not heteroplasmic (binomial test for departure from equal frequency *p* = 2.2 × 10^−16^ in all cases). The two Gymnosperms in our dataset both contain only haplotype B, and one of the Pteridophytes in our dataset (*Selaginella tamariscina*) contains only haplotype C. Fig. 2 shows clearly that, as expected, there is a trend for datasets with larger sample sizes to show relative haplotype abundances closer to exactly 50/50.

**Table 1.**
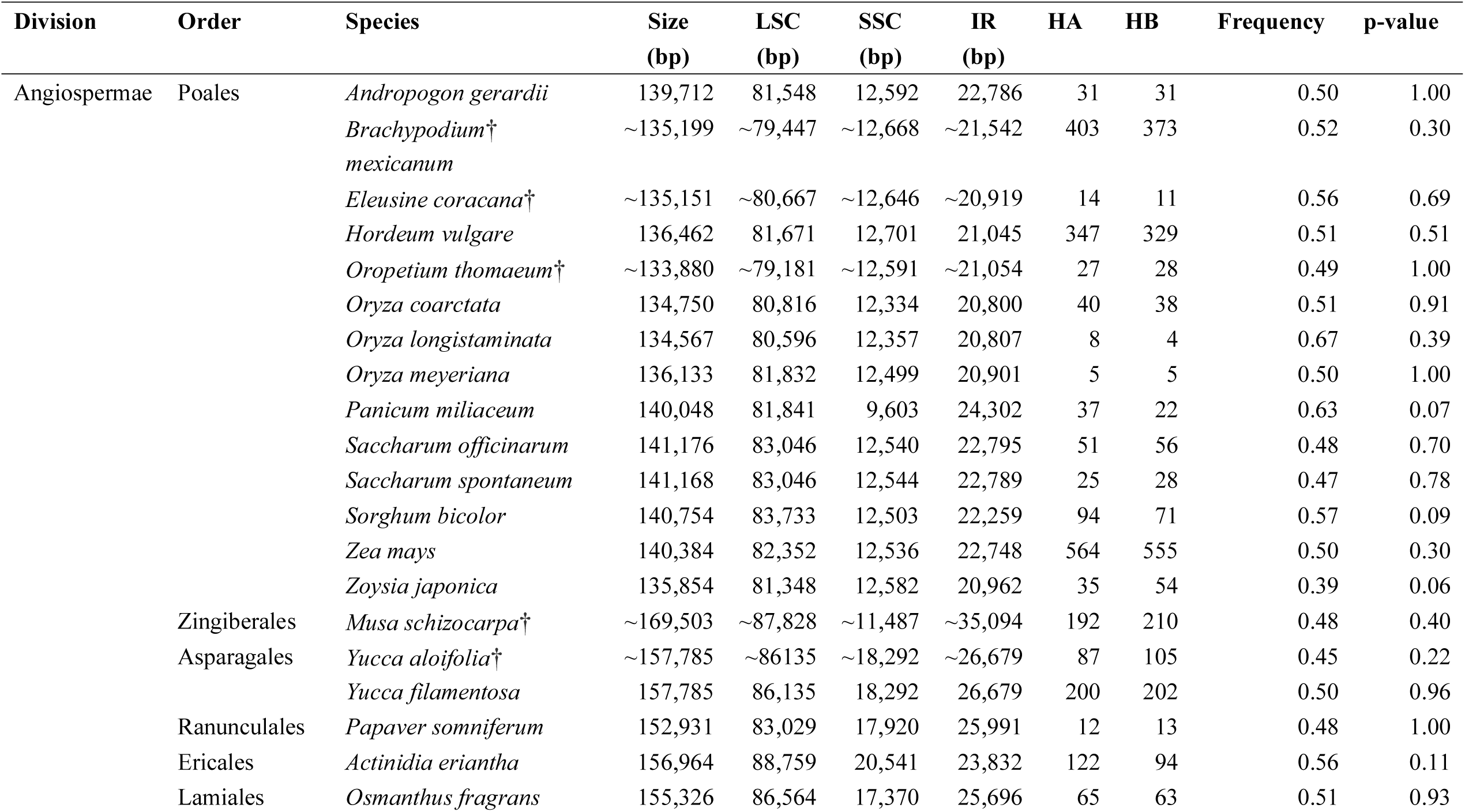
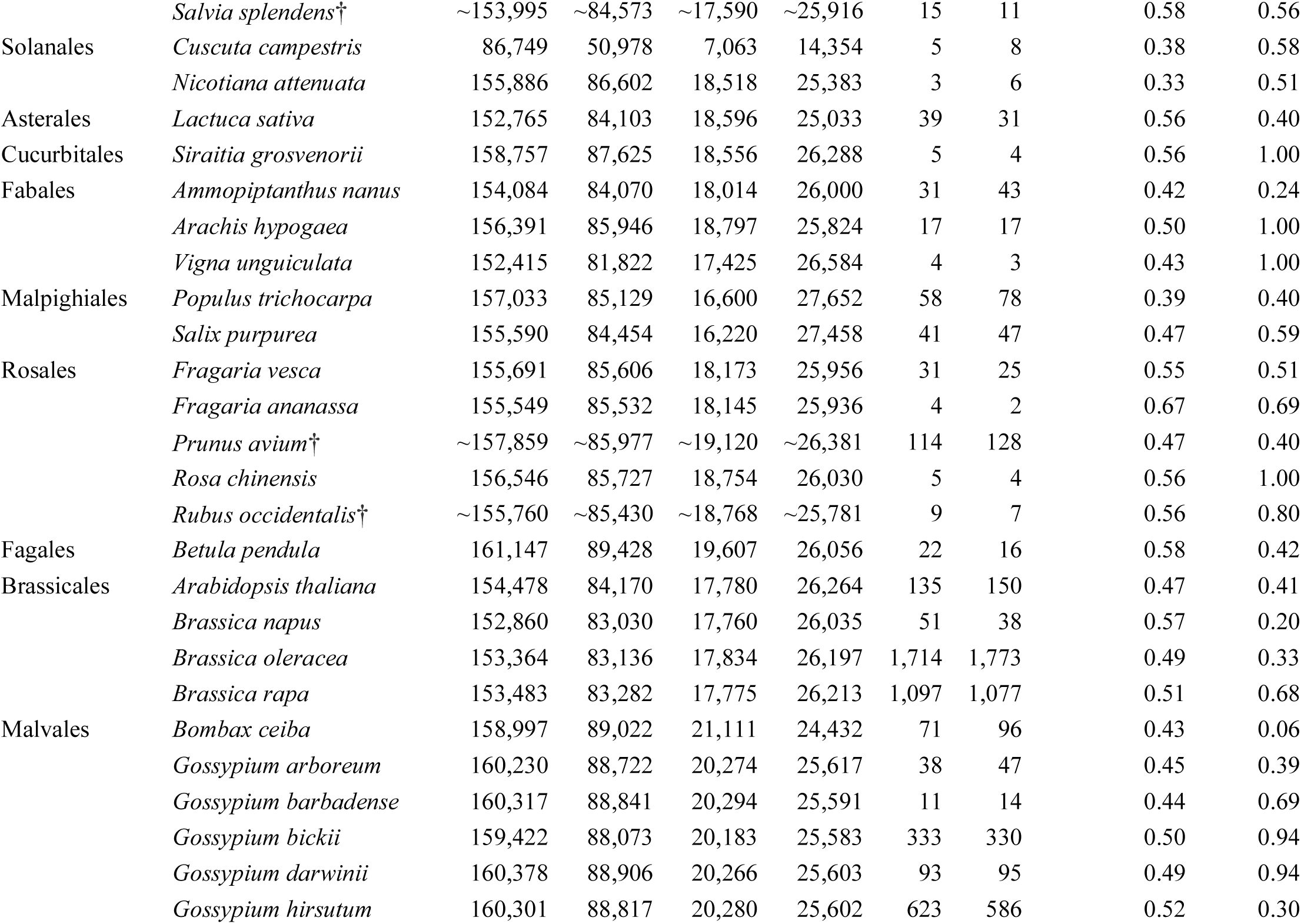
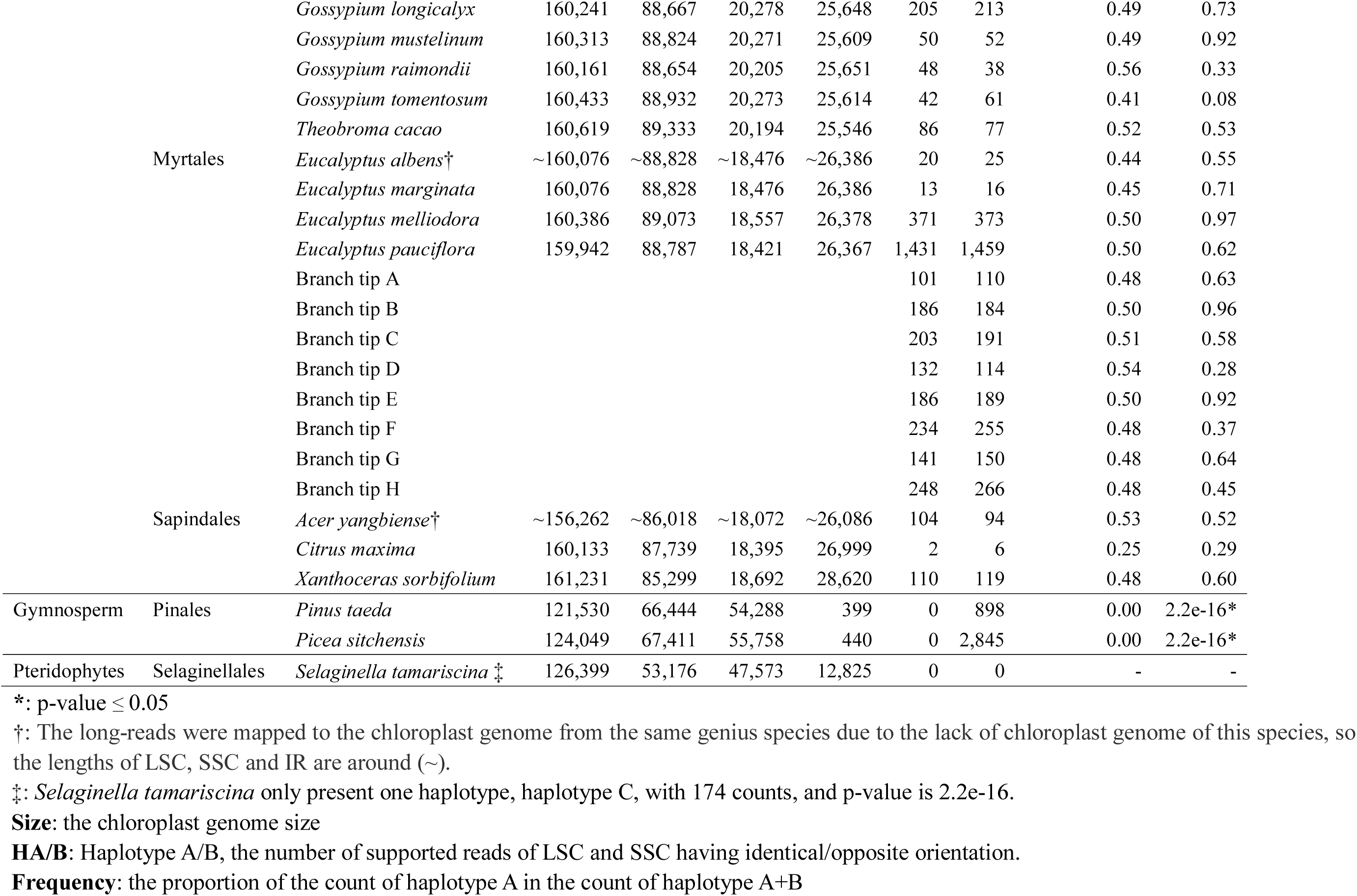
The frequency of existing chloroplast genome haplotypes from 61 species.

| Division | Order | Species | Size<br>(bp) | LSC<br>(bp) | SSC<br>(bp) | IR<br>(bp) | HA | HB | Frequency | p-value |
| --- | --- | --- | --- | --- | --- | --- | --- | --- | --- | --- |
| Angiospermae | Poales | *Andropogon gerardii* | 139,712 | 81,548 | 12,592 | 22,786 | 31 | 31 | 0.50 | 1.00 |
|  |  | *Brachypodium*†<br>*mexicanum* | ~135,199 | ~79,447 | ~12,668 | ~21,542 | 403 | 373 | 0.52 | 0.30 |
|  |  | *Eleusine coracana*† | ~135,151 | ~80,667 | ~12,646 | ~20,919 | 14 | 11 | 0.56 | 0.69 |
|  |  | *Hordeum vulgare* | 136,462 | 81,671 | 12,701 | 21,045 | 347 | 329 | 0.51 | 0.51 |
|  |  | *Oropetium thomaeum*† | ~133,880 | ~79,181 | ~12,591 | ~21,054 | 27 | 28 | 0.49 | 1.00 |
|  |  | *Oryza coarctata* | 134,750 | 80,816 | 12,334 | 20,800 | 40 | 38 | 0.51 | 0.91 |
|  |  | *Oryza longistaminata* | 134,567 | 80,596 | 12,357 | 20,807 | 8 | 4 | 0.67 | 0.39 |
|  |  | *Oryza meyeriana* | 136,133 | 81,832 | 12,499 | 20,901 | 5 | 5 | 0.50 | 1.00 |
|  |  | *Panicum miliaceum* | 140,048 | 81,841 | 9,603 | 24,302 | 37 | 22 | 0.63 | 0.07 |
|  |  | *Saccharum officinarum* | 141,176 | 83,046 | 12,540 | 22,795 | 51 | 56 | 0.48 | 0.70 |
|  |  | *Saccharum spontaneum* | 141,168 | 83,046 | 12,544 | 22,789 | 25 | 28 | 0.47 | 0.78 |
|  |  | *Sorghum bicolor* | 140,754 | 83,733 | 12,503 | 22,259 | 94 | 71 | 0.57 | 0.09 |
|  |  | *Zea mays* | 140,384 | 82,352 | 12,536 | 22,748 | 564 | 555 | 0.50 | 0.30 |
|  |  | *Zoysia japonica* | 135,854 | 81,348 | 12,582 | 20,962 | 35 | 54 | 0.39 | 0.06 |
|  | Zingiberales | *Musa schizocarpa*† | ~169,503 | ~87,828 | ~11,487 | ~35,094 | 192 | 210 | 0.48 | 0.40 |
|  | Asparagales | *Yucca aloifolia*† | ~157,785 | ~86,135 | ~18,292 | ~26,679 | 87 | 105 | 0.45 | 0.22 |
|  |  | *Yucca filamentosa* | 157,785 | 86,135 | 18,292 | 26,679 | 200 | 202 | 0.50 | 0.96 |
|  | Ranunculales | *Papaver somniferum* | 152,931 | 83,029 | 17,920 | 25,991 | 12 | 13 | 0.48 | 1.00 |
|  | Ericales | *Actinidia eriantha* | 156,964 | 88,759 | 20,541 | 23,832 | 122 | 94 | 0.56 | 0.11 |
|  | Lamiales | *Osmanthus fragrans* | 155,326 | 86,564 | 17,370 | 25,696 | 65 | 63 | 0.51 | 0.93 |

|  |  |  |  |  |  |  |  |  |  |
| --- | --- | --- | --- | --- | --- | --- | --- | --- | --- |
|  | *Salvia splendens*† | ~153,995 | ~84,573 | ~17,590 | ~25,916 | 15 | 11 | 0.58 | 0.56 |
| Solanales | *Cuscuta campestris* | 86,749 | 50,978 | 7,063 | 14,354 | 5 | 8 | 0.38 | 0.58 |
|  | *Nicotiana attenuata* | 155,886 | 86,602 | 18,518 | 25,383 | 3 | 6 | 0.33 | 0.51 |
| Asterales | *Lactuca sativa* | 152,765 | 84,103 | 18,596 | 25,033 | 39 | 31 | 0.56 | 0.40 |
| Cucurbitales | *Siraitia grosvenorii* | 158,757 | 87,625 | 18,556 | 26,288 | 5 | 4 | 0.56 | 1.00 |
| Fabales | *Ammopiptanthus nanus* | 154,084 | 84,070 | 18,014 | 26,000 | 31 | 43 | 0.42 | 0.24 |
|  | *Arachis hypogaea* | 156,391 | 85,946 | 18,797 | 25,824 | 17 | 17 | 0.50 | 1.00 |
|  | *Vigna unguiculata* | 152,415 | 81,822 | 17,425 | 26,584 | 4 | 3 | 0.43 | 1.00 |
| Malpighiales | *Populus trichocarpa* | 157,033 | 85,129 | 16,600 | 27,652 | 58 | 78 | 0.39 | 0.40 |
|  | *Salix purpurea* | 155,590 | 84,454 | 16,220 | 27,458 | 41 | 47 | 0.47 | 0.59 |
| Rosales | *Fragaria vesca* | 155,691 | 85,606 | 18,173 | 25,956 | 31 | 25 | 0.55 | 0.51 |
|  | *Fragaria ananassa* | 155,549 | 85,532 | 18,145 | 25,936 | 4 | 2 | 0.67 | 0.69 |
|  | *Prunus avium*† | ~157,859 | ~85,977 | ~19,120 | ~26,381 | 114 | 128 | 0.47 | 0.40 |
|  | *Rosa chinensis* | 156,546 | 85,727 | 18,754 | 26,030 | 5 | 4 | 0.56 | 1.00 |
|  | *Rubus occidentalis*† | ~155,760 | ~85,430 | ~18,768 | ~25,781 | 9 | 7 | 0.56 | 0.80 |
| Fagales | *Betula pendula* | 161,147 | 89,428 | 19,607 | 26,056 | 22 | 16 | 0.58 | 0.42 |
| Brassicales | *Arabidopsis thaliana* | 154,478 | 84,170 | 17,780 | 26,264 | 135 | 150 | 0.47 | 0.41 |
|  | *Brassica napus* | 152,860 | 83,030 | 17,760 | 26,035 | 51 | 38 | 0.57 | 0.20 |
|  | *Brassica oleracea* | 153,364 | 83,136 | 17,834 | 26,197 | 1,714 | 1,773 | 0.49 | 0.33 |
|  | *Brassica rapa* | 153,483 | 83,282 | 17,775 | 26,213 | 1,097 | 1,077 | 0.51 | 0.68 |
| Malvales | *Bombax ceiba* | 158,997 | 89,022 | 21,111 | 24,432 | 71 | 96 | 0.43 | 0.06 |
|  | *Gossypium arboreum* | 160,230 | 88,722 | 20,274 | 25,617 | 38 | 47 | 0.45 | 0.39 |
|  | *Gossypium barbadense* | 160,317 | 88,841 | 20,294 | 25,591 | 11 | 14 | 0.44 | 0.69 |
|  | *Gossypium bickii* | 159,422 | 88,073 | 20,183 | 25,583 | 333 | 330 | 0.50 | 0.94 |
|  | *Gossypium darwinii* | 160,378 | 88,906 | 20,266 | 25,603 | 93 | 95 | 0.49 | 0.94 |
|  | *Gossypium hirsutum* | 160,301 | 88,817 | 20,280 | 25,602 | 623 | 586 | 0.52 | 0.30 |
|  |  | *Gossypium longicalyx* | 160,241 | 88,667 | 20,278 | 25,648 | 205 | 213 | 0.49 | 0.73 |
|  |  | *Gossypium mustelinum* | 160,313 | 88,824 | 20,271 | 25,609 | 50 | 52 | 0.49 | 0.92 |
|  |  | *Gossypium raimondii* | 160,161 | 88,654 | 20,205 | 25,651 | 48 | 38 | 0.56 | 0.33 |
|  |  | *Gossypium tomentosum* | 160,433 | 88,932 | 20,273 | 25,614 | 42 | 61 | 0.41 | 0.08 |
|  |  | *Theobroma cacao* | 160,619 | 89,333 | 20,194 | 25,546 | 86 | 77 | 0.52 | 0.53 |
|  | Myrtales | *Eucalyptus albens*† | ~160,076 | ~88,828 | ~18,476 | ~26,386 | 20 | 25 | 0.44 | 0.55 |
|  |  | *Eucalyptus marginata* | 160,076 | 88,828 | 18,476 | 26,386 | 13 | 16 | 0.45 | 0.71 |
|  |  | *Eucalyptus melliodora* | 160,386 | 89,073 | 18,557 | 26,378 | 371 | 373 | 0.50 | 0.97 |
|  |  | *Eucalyptus pauciflora* | 159,942 | 88,787 | 18,421 | 26,367 | 1,431 | 1,459 | 0.50 | 0.62 |
|  |  | Branch tip A |  |  |  |  | 101 | 110 | 0.48 | 0.63 |
|  |  | Branch tip B |  |  |  |  | 186 | 184 | 0.50 | 0.96 |
|  |  | Branch tip C |  |  |  |  | 203 | 191 | 0.51 | 0.58 |
|  |  | Branch tip D |  |  |  |  | 132 | 114 | 0.54 | 0.28 |
|  |  | Branch tip E |  |  |  |  | 186 | 189 | 0.50 | 0.92 |
|  |  | Branch tip F |  |  |  |  | 234 | 255 | 0.48 | 0.37 |
|  |  | Branch tip G |  |  |  |  | 141 | 150 | 0.48 | 0.64 |
|  |  | Branch tip H |  |  |  |  | 248 | 266 | 0.48 | 0.45 |
|  | Sapindales | *Acer yangbiense*† | ~156,262 | ~86,018 | ~18,072 | ~26,086 | 104 | 94 | 0.53 | 0.52 |
|  |  | *Citrus maxima* | 160,133 | 87,739 | 18,395 | 26,999 | 2 | 6 | 0.25 | 0.29 |
|  |  | *Xanthoceras sorbifolium* | 161,231 | 85,299 | 18,692 | 28,620 | 110 | 119 | 0.48 | 0.60 |
| Gymnosperm | Pinales | *Pinus taeda* | 121,530 | 66,444 | 54,288 | 399 | 0 | 898 | 0.00 | 2.2e-16* |
|  |  | *Picea sitchensis* | 124,049 | 67,411 | 55,758 | 440 | 0 | 2,845 | 0.00 | 2.2e-16* |
| Pteridophytes | Selaginellales | *Selaginella tamariscina* ‡ | 126,399 | 53,176 | 47,573 | 12,825 | 0 | 0 | - | - |

\*: p-value $\leq 0.05$
†: The long-reads were mapped to the chloroplast genome from the same genus species due to the lack of chloroplast genome of this species, so the lengths of LSC, SSC and IR are around (~).
‡: *Selaginella tamariscina* only present one haplotype, haplotype C, with 174 counts, and p-value is 2.2e-16. **Size:** the chloroplast genome size **HA/B:** Haplotype A/B, the number of supported reads of LSC and SSC having identical/opposite orientation. **Frequency:** the proportion of the count of haplotype A in the count of haplotype A+B

**Fig. 1.**
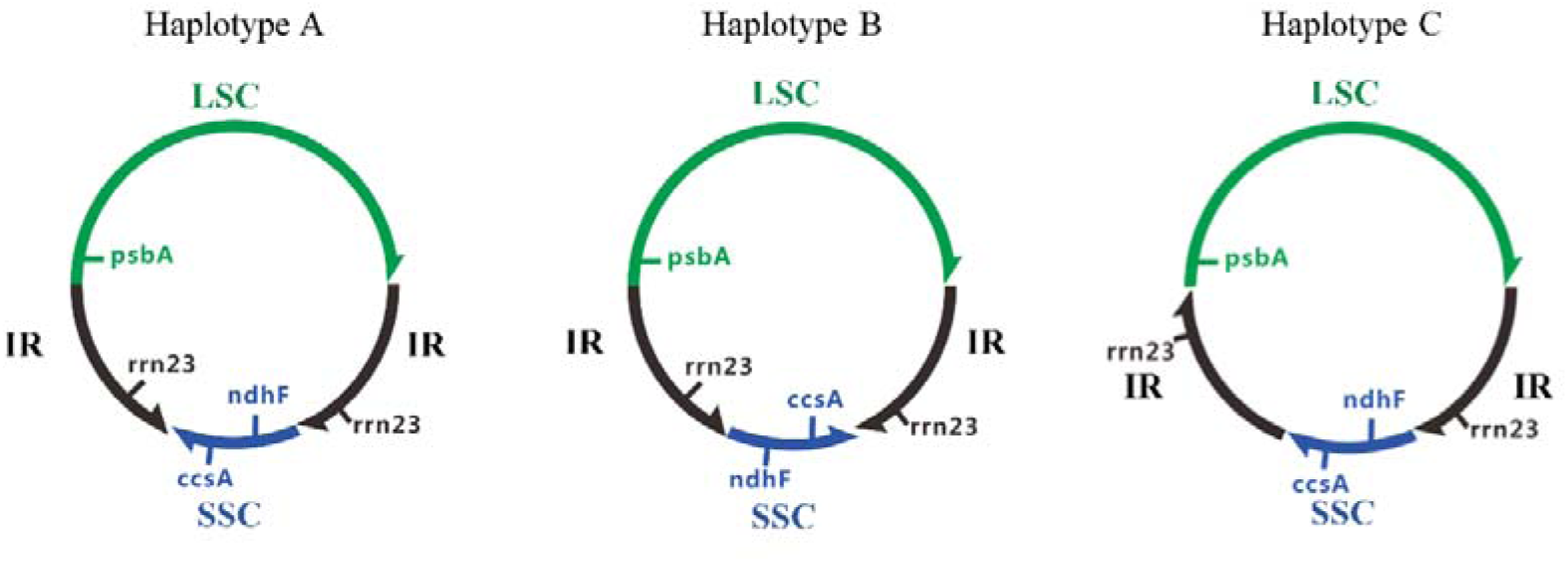
The three different structural haplotypes of chloroplast genomes detected in this study. The green region is the Long Single Copy (LSC) region and blue region is the Short Single Copy (SSC) region. The two black regions are the Inverted Repeat (IR) regions. The arrow denotes 5’-3’ orientation. *psbA* is in the minus strand of LSC region, whereas *rrn23* is in the plus strand of IR regions. *ndhF* is in the minus strand of SSC region, while *ccsA* is in the plus strand of SSC region. For ease of communication, we use the relative order of three genes (*psbA* in LSC, and *ndhF* and *ccsA* in the SSC) to label these two haplotypes ‘A’ and ‘B’. In haplotype A, these genes are ordered *psbA*-*ndhF*-*ccsA*. In haplotype B these genes are ordered *psbA*-*ccsA*-*ndhF*. For haplotype C, these three genes are ordered the same as haplotype A, but the repeat regions are in-line rather than inverted.

**Fig. 2.**
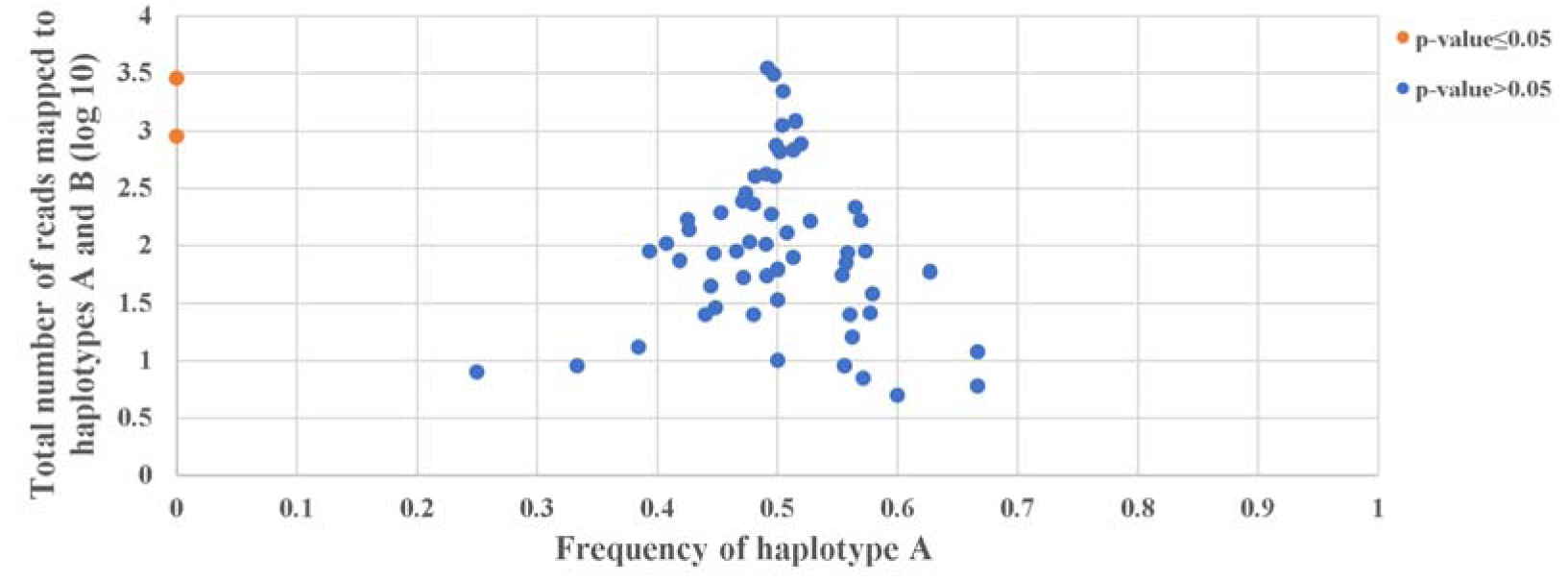
The relationship between haplotype counts and haplotype frequencies. Each point is one species. A blue dot means binomial p-value > 0.05, whereas the orange dot means binomial p-value ≤ 0.05. Two species, *Pinus taeda, Picea sitchensis* only showed evidence of haplotype B, therefore the frequencies of them were 0. For the remaining 58 species, they show higher level of equal frequency between haplotype A and B with the increase of sample size. One species (*Selaginella tamariscina*) is omitted from this figure because it contained only a third haplotype (haplotype C, Figure 1).

Across 8 different samples taken from a single large *Eucaltyptus pauciflora* plant (Table 1), we detected roughly equal frequencies of haplotypes A and B in all 8 samples (binomial p-values > 0.05 in all cases), no evidence for differences in relative frequencies among samples (chi-square p-value > 0.05), and no evidence for phylogenetic signal in the any of the relative abundances (p-value > 0.05 in all tests), where the underlying topology and branch lengths are taken to be the physical structure of the tree itself (see Materials and Methods).

## Discussion

In this study, we develop a new method to detect and quantify structural heteroplasmies in the chloroplast genome. By applying this method to a broad range of plant species including representatives from Angiosperms, Gymnosperms and Pteridophytes, we show that of the large number of possible structural haplotypes, only a very small number are observed in nature. For example, species with IRs almost universally contain the same two structural haplotypes in ratios that do not depart significantly from 50/50. The only exceptions to this are species in which the repeats are not inverted (e.g. one of the Pteridophytes in our dataset), or in which the repeats are inverted but much reduced in length (e.g. both of the Gymnosperms in our dataset). In these species, we detect just one haplotype. Our results demonstrate not only that the relative abundance of the two most common haplotypes stable across the Angiosperm tree of life (i.e. all 58 Angiosperms in our dataset had a frequency that was not different from 50/50), but that it also stable within an individual plant (i.e. within a single large plant, all sampled leaves contained both haplotypes at a ratio that was not different from 50/50). This suggests that the process which maintains the equal frequencies of the two commonly-observed haplotypes operates relatively rapidly.

The most parsimonious interpretation of our results suggests that structural heteroplamsy should be extremely common across all angiosperms. In light of this, it may be necessary to re-examine many previous suggestions of structural differences in the chloroplast genomes of angiosperms *between* species (9-13), as Emery *et al*(14) suggested. For example, Ibrahim *et al* (9) implied that the SSC of *Gossypium barbadense* was inverted when compared to *Gossypium hirsutum*. However, our study shows that both *Gossypium barbadense* and *Gossypium hirsutum* contain two structural haplotypes (haplotypes A and B (Table 1)), thus showing that there are no structural differences between these two species. Future studies may benefit from assuming at the outset that it is highly likely that the chloroplast genome of any angiosperm species, particularly if it contains two long inverted repeats, is most likely to exist in two roughly equimolar haplotypes, haplotypes A and B shown in Fig. 1. This phenomenon could also exist in Gymnosperms and Pteridophytes, but in this study we don’t have enough available data to provide solid evidence to support this.

Of the 61 species for which we had sufficient data, we find just three species in which there is no evidence for structural heteroplasmy in the chloroplast genome: *Selaginella tamariscina, Pinus taeda* and *Picea sitchensis. Selaginella tamariscina* is a pteridophyte in which the repeats in the chloroplast genome are positioned in-line rather than inverted (20). The existence of a single structural haplotype in this species is therefore consistent with the hypothesis that the process which generates chloroplast structural heteroplasmy is mediated by long IRs. In the gymnosperms *Pinus taeda* and *Picea sitchensis*, the repeats are inverted but their lengths are highly reduced from the typical 10-30 kb in most species to just 399 bp and 440 bp, respectively. The existence of a single structural haplotype in these species is also consistent with the flip-flop recombination theory, because it is feasible that the highly reduced IRs preclude the formation of the dumbbell-like structure (4) which is necessary to activate flip-flop recombination. Whether or not flip-flop recombination is the underlying cause, our results confirm that chloroplast structural heteroplasmy appears to require the existence of two long IRs in the chloroplast genome, and therefore that groups which lack this trait would be expected to lack chloroplast structural heteroplasmy. For example, chloroplast genomes in the Pinaceae and Cupressophytes (two major groups of gymnosperms) usually lack one IR or have IRs that are highly reduced in size (15, 21-26), and we therefore expect that these species would lack chloroplast structural heteroplasmy.

In this study, we focussed on examining heteroplasmies in the quadripartite structure of the chloroplast genome. However, other smaller structural heteroplasmies have been observed in some chloroplast genomes, and their existence challenges the conclusion that long inverted repeats are a pre-requisite for the existence of structural heteroplasmy in the more general sense. Although flip-flop recombination may be absent when the IR regions are small (see above), some studies have suggested that homologous recombination could induce structural heterplasmy from other much smaller inverted repeats in chloroplast genomes (25, 26). For instance, the *trnQ-UUG* gene (∼150 bp) is duplicated in the LSC region in some *Juniperus* species, and this duplication is associated with a 36 kb inversion resulting in the coexistence of two isomeric chloroplast genome structures, although their abundance is unequal (0.8% - 5.0%) (25). Qu *et al* (26) reported a similar finding in other Cupressoideae species, and suggest that even a pair of very small inverted repeats (11 bp) could induce a 34 kb inversion and resulting structural heteroplasmy in *Calocedrus macrolepis*. In light of these results, it remains unclear why the highly reduced IRs of *Pinus taeda* and *Picea sitchensis* studied here fail to induce structural heteroplasmy, while much shorter inverted repeats in regions of the chloroplast genome outside the IR region apparently do induce structural heteroplasmy. One possibility is that different processes mediate the formation of structural heteroplasmies in these cases (25, 26). Another possibility is that the frequency of the alternative haplotype is too low for us to detect in *Pinus taeda* and *Picea sitchensis.* However, if this were the case the frequency of haplotype A in these species would have to be extremely low: for example in *Picea sitchensis* we detected 2845 long reads supporting haplotype B and no reads supporting haplotype A.

The Cp-hap pipeline we present here provides a convenient method for quantifying structural heteroplasmy in chloroplast genomes. It adds to suite of existing methods such as restriction digests (3, 4) and BES (5), but differs from these methods in that it uses long sequencing reads to provide direct evidence for uniquely-identifiable chloroplast structures. It is of note that the two available long-read sequencing methods, PacBio and Oxford Nanopore sequencing, differ in their utility for detecting chloroplast structural heteroplasmy. For example, only ∼63% (46 out of 73 species) of the PacBio datasets we used in this study contained enough long chloroplast reads to adequately measure chloroplast structural heteroplasmy. In contrast, ∼88% (15 out of 17 species) of Oxford Nanopore datasets provided enough long-read to measure the chloroplast genome heteroplasmy. Given the increasingly popularity of Oxford Nanopore sequencing, and ongoing improvements in the read-lengths available from PacBio sequencers, we hope that the simplicity of the Cp-hap pipeline will accelerate further work on chloroplast structural heteroplasmy.

## Materials and Methods

### Cp-hap, a new method for quantifying chloroplast structural haplotypes using long-read sequencing data

New long-read sequencing methods such as those developed by Oxford Nanopore can routinely sequence single DNA molecules of 10s of kb in length. These molecules can be used to provide direct evidence for the existence and relative abundance of a large range of chloroplast genome haplotypes. Furthermore, long-reads from the chloroplast genome are often highly abundant in plant genome sequencing projects, because the chloroplast genome is typically present in copy numbers at least two orders of magnitude higher than the nuclear genome (27, 28). Thus, most whole genome sequencing projects of plant provide abundant information for quantifying chloroplast structural haplotypes. Here, we describe a simple approach that uses this long-read sequencing data to estimate and quantify chloroplast structural haplotypes. We call this approach the Cp-hap pipeline, and provide open source code and detailed instructions for running this pipeline on GitHub at https://github.com/asdcid/Cp-hap. The Cp-hap pipeline consists of three steps: (i) create a fasta file that contains all haplotypes of interest; (ii) map the long-reads to the potential structures in step (i); (iii) use the results of step (ii) to quantify the relative abundance of all possible haplotypes. Below, we describe each of these steps in more detail.

Step (i) requires us to create a fasta file containing all of the structural haplotypes of interest. To create a list of all possible haplotypes from the LSC, SSC, and IR regions, we first consider that each of these regions can have four different orientations: original, reversed, complement, and reverse-complement (Fig. S3A). This results in 256 possible structural haplotypes (256 = 4×4×4×4) in which the regions retain the ordering conserved across plant chloroplast genomes of LSC-IRA-SSC-IRB. These 256 haplotypes can be grouped into 128 identifiable structures, since it is not possible to distinguish between one structure and its direct complement given the two-stranded nature of the DNA molecule.

To uniquely identify one of these 128 structures, a single sequencing read would need to cover at least some parts of all four regions (LSC, SSC and the two IR regions), for which the read would need to be at least 30-50 kb. This is because to cover all four regions, at a minimum a read must entirely cover the SSC (∼20 kb) region and one IR region (10-30 kb) and at least partially cover the LSC region and the other IR region. But the abundance of reads of this length tends to be relatively low in most datasets, meaning that in most cases it is almost impossible to uniquely identify one of the 128 possible structures by simply mapping a read to each structure.

To reduce the length of long-reads required to uniquely identify chloroplast structural haplotypes, the Cp-hap pipeline assumes by default that the two large repeat regions are always inverted. When assuming that the IR regions are always inverted, there are only 32 uniquely identifiable chloroplast genome structural haplotypes (Fig. S1). In this situation, a read only needs to entirely cover one IR region and partially cover the two adjacent LSC and SSC regions to provide evidence to uniquely identify one of the 32 structures (Fig. S3B). Because of this, this approach can provide direct evidence for one of the 32 possible structures using reads that are just 10-30 kb in size (i.e. at least as long as the length of a single IR region). Reads of this length are often highly abundant in long-read datasets, meaning that the default Cp-hap method is readily applicable to many long-read datasets being produced today.

In Step (i), the default Cp-hap pipeline creates a fasta file of the 32 possible structural haplotypes when assuming that the IRs are inverted by simply combining the LSC, SSC and IR region sequences with different orientations. After creating each structural haplotype, we then duplicate and concatenate it, such that single reads which span the point at which the genome was linearised will still align successfully to the relevant haplotype in the fasta file. This creates a fasta file with 32 sequences, each of which represents one of the 32 uniquely identifiable chloroplast structural haplotypes, and each of which contains a sequence that is twice as long as the chloroplast genome of interest.

In step (ii) of the pipeline, we align our long-reads to our reference set of 32 possible structural haplotypes with Minimap2 (29), using “–secondary=no” setting to ensure that only the best alignment for each read is retained. After aligning all long-reads to the set of 32 possible structural haplotypes, we examine each mapped read, and retain only those that completely cover a whole IR region and at least 1 kb of the adjacent LSC and SSC regions. We then consider that these reads, which we term valid reads, uniquely identify one of the 32 possible structural haplotypes.

In step (iii) of the Cp-hap pipeline, we calculate the relative proportion of each of the 32 possible haplotypes. To do this, we simply divide the number of valid reads that aligned to each of the haplotypes by total number of reads that aligned to all haplotypes. These proportions provide direct estimates of the relative abundance of all 32 uniquely identifiable chloroplast genome structural haplotypes.

However, we acknowledge that the IR regions are not always the case. For example, the large repeat regions are positioned in-line instead of inverted in *Selaginella tamariscina* chloroplast genome (20). In cases such as this, Cp-hap pipeline is also able to figure out the structure. Here, we provide an example that how Cp-hap pipeline confirms a chloroplast genome with atypical structure, such as a paired of in-line repeats (Fig. S3C). Since Cp-hap duplicates and concatenates all default 32 structures (with IRs), reads from chloroplast genome with in-line repeats are able to map to Block A and/or C region of LSC_IR_SSC_IRrc structure (one of the 32 default structure in Cp-hap pipeline), or Block B region of LSC_IRrc_SSC_IR structure (the other default structure in Cp-hap pipeline). Those reads are impossible to map to Block B region of LSC_IR_SSC_IRrc, or Block A or C region of LSC_IRrc_SSC_IR. Therefore, if some reads only map to Block A and/or C region of LSC_IR_SSC_IRrc and Block B region of LSC_IRrc_SSC_IR, it suggests that the chloroplast genome acquires in-line repeats.

In sum, Cp-hap pipeline can provide direct evidence to identify one of the 32 chloroplast structural haplotypes with a pair of IRs, and provide indirect evidence to identify other 96 chloroplast genome structural haplotypes with a pair of non-inverted repeats.

### Assessing chloroplast structural heteroplasmy across the plant tree of life

We used the Cp-hap pipeline to assess chloroplast genome structural heteroplasmy across the plant tree of life. To do this we searched for publicly available long-read data from land plants in the NCBI SRA database (https://www.ncbi.nlm.nih.gov). We only selected datasets in which the average read length was >9 kb and for which the total amount of sequence data was >5Gb. 90 of the 164 potential species with datasets met this requirement in April, 2019.

For the 90 species, 19 of them do not have available chloroplast reference genomes. Due to the relative conservation of land plant chloroplast genomes, we used the chloroplast genome in the same genus as the reference genome for 17 species (Dataset S1). For the final two species, *Herrania umbratica* and *Siraitia grosvenorii*, we assembled and annotated chloroplast genomes *de novo* using long and short-reads (Dataset S1) following the pipeline described in (30). The chloroplast genome of *Herrania umbratica* and *Siraitia grosvenorii* have been depostied in NCBI (MN163033 and MK279915).

Full details of the reads, accession numbers, and chloroplast genomes used for each of the 90 species in our dataset are given in Dataset S1.

### Chloroplast structural haplotype quantification

For the 90 species for which we have long-read data, we ran the Cp-hap pipeline using default settings to generate the 32 possible haplotypes from the LSC, SSC, and IR regions. We then ran the rest of the Cp-hap pipeline with default settings for all species, resulting in measurements for each species of the number of valid reads that mapped to each haplotype, and the relative abundance of each haplotype. We calculated the relative abundance of each haplotype only if the species had more than five valid reads in total. This filter led to us excluding 29 datasets that did not have more than five valid chloroplast reads, resulting in a final dataset of 61 species for which we have sufficient long-read data to estimate relative haplotype abundance.

For the *Eucalyptus pauciflora* dataset, we quantified the chloroplast structural haplotypes not only for a single individual, but also for eight separate samples taken from different branches of the same individual. This was possible because this data was sequenced from leaves collected from eight different branch tips from a single individual plant, and each sample contained sufficient long chloroplast reads to provide reliable haplotype quantification. This provides a unique opportunity to understand whether the relative abundance of chloroplast structural haplotypes varies or remains constant within an individual, potentially providing insights into the mechanisms which lead to the existence of multiple haplotypes.

### Statistical analyses

Although our method can detect a large number of structural haplotypes, across all of the species we examined with inverted repeats, we observed three haplotypes, haplotypes A, B, and C (Fig. 1).

Because no species contained more than two haplotypes, we used a binomial test to ask whether the abundances of haplotype A and B differ from a 1:1 ratio in each species, or in each individual sample in the case of *Eucalyptus pauciflora*. A significant result from a binomial test suggests that the observed haplotype abundances are unlikely to be explained by an underlying 1:1 ratio of chloroplast structural haplotypes.

In addition, since we have data for eight different branch tips of a single *Eucalyptus pauciflora* individual, we used two additional approaches to test for differences between samples. First, we used a chi-square test to ask whether the abundances of the two haplotypes differ among samples. Second, we tested for phylogenetic signal among the relative abundance of haplotypes observed in the eight branch tips. To do this, we used the structure of the physical tree (with branch lengths in units of centimetres) and the phylosignal package (31) in R (32). This package implements five commonly-used tests of phylogenetic signal: (i) Abouheif’s Cmean (33); (ii & iii) Blomberg’s K and K* (34); (iv) Moran’s I (35); (v) Pagel’s Lambda (36). These tests use different approaches to ask whether the physical structure of the individual tree helps to explain the observed variation in the relative haplotype abundance of the eight samples. For example, if the process that generates different haplotypes operates relatively slowly with respect to the age of this individual, which is roughly 40 years, then we might expect haplotype frequencies to vary due to genetic drift, and thus for the haplotype frequencies in neighbouring parts of the tree to be more similar than would be expected by chance. This similarity would be revealed by the detection of significant phylogenetic signal in the relative abundance data. However, if the process that generates the different haplotypes operates very quickly with respect to the age of this individual, then we would expect all haplotype frequencies to be roughly equal, and thus we would expect to observe no phylogenetic signal in the haplotype abundance data.

## Acknowledgments

We thank the help of Chen Han in the statistical analysis. We also thank the people below for providing data: Xuzhi Chao (for *Selaginella tamariscina* data), Hongyan Du (for *Eucommia ulmoides* data). This work was supported by grants from the Australian Research Council Future Fellowship, FT140100843.

