## Supplementary figures and images for "Stable and widespread structural heteroplasmy in chloroplast genomes revealed by a new long-read quantification method"

### Fig. S1

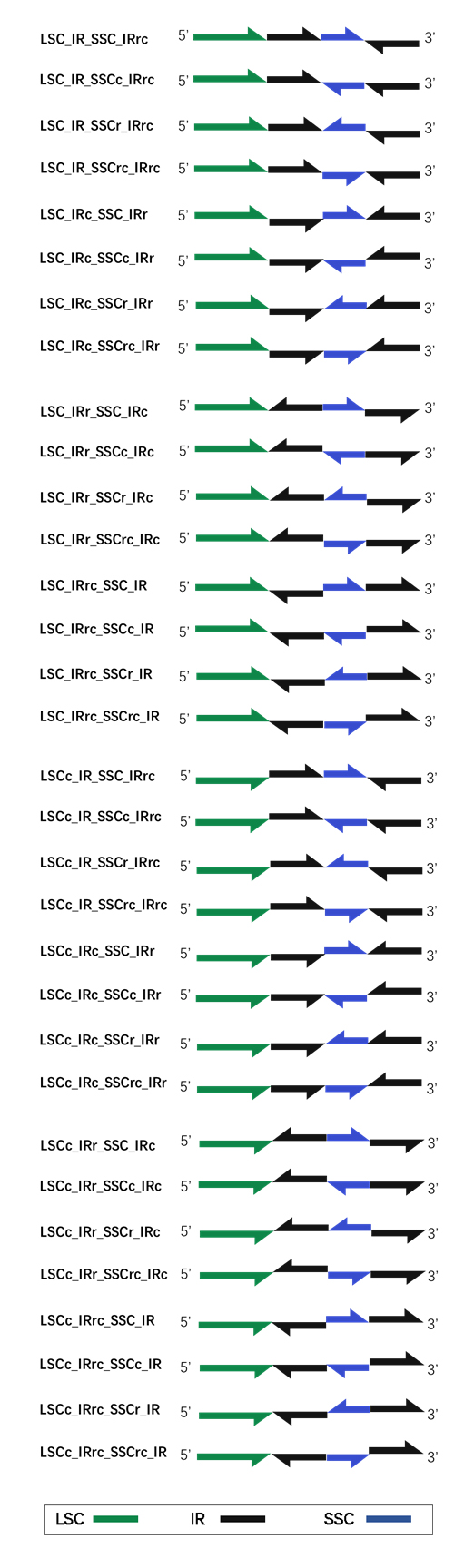

### Fig. S2

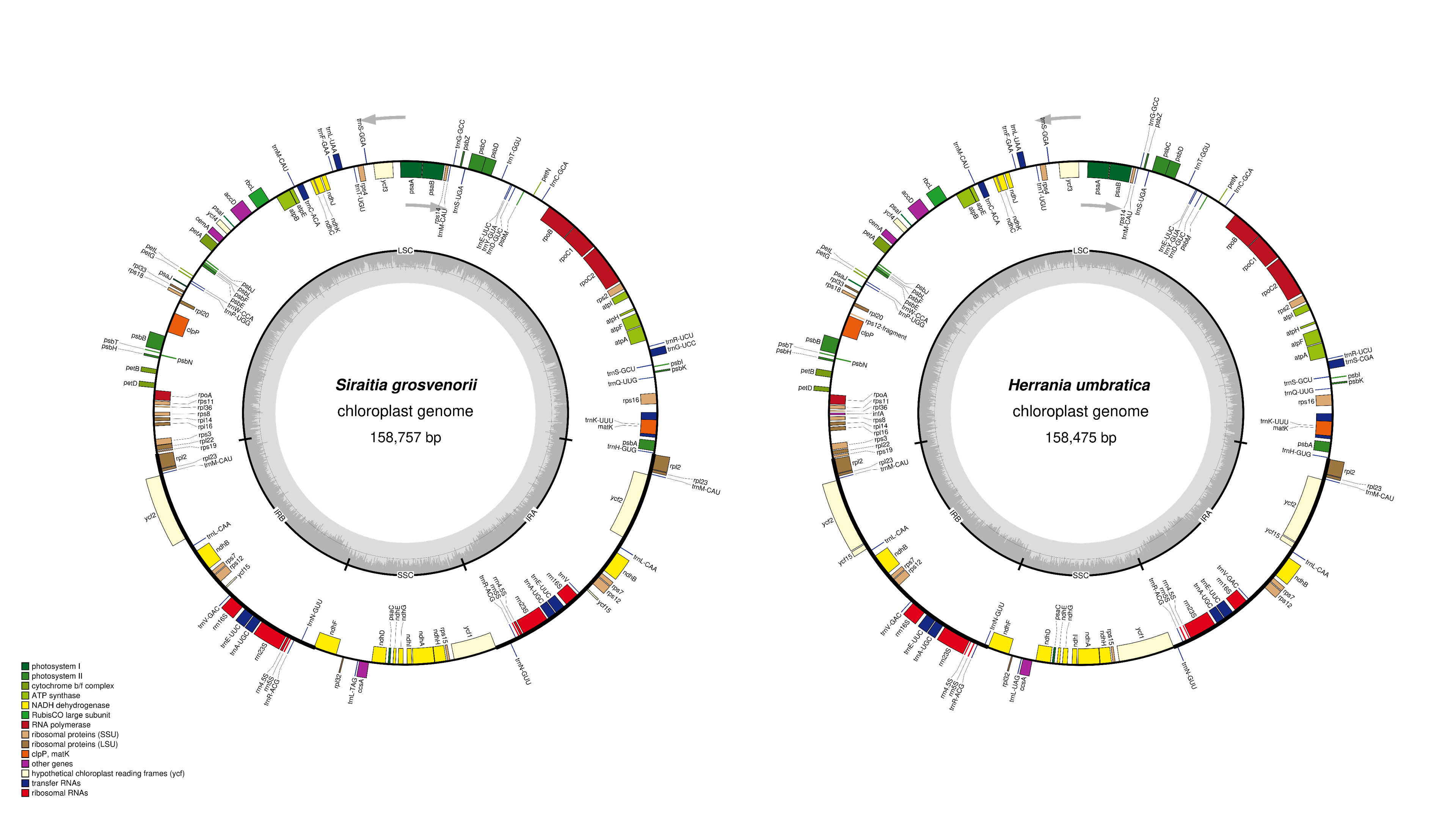

### Fig. S3

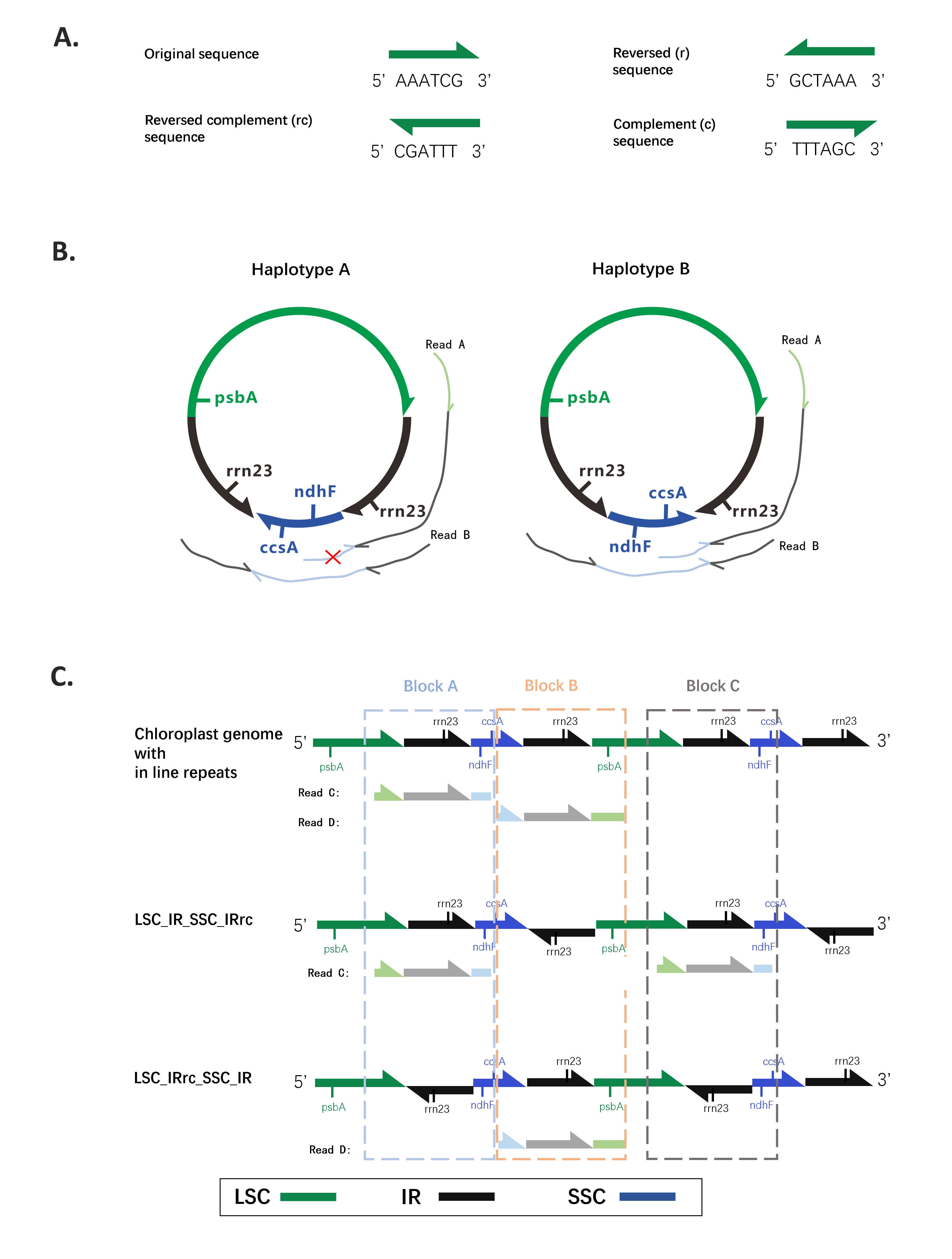
