## Supplementary material for "Stable and widespread structural heteroplasmy in chloroplast genomes revealed by a new long-read quantification method": Table S2

**Table S2**. The frequency of existing chloroplast genome haplotypes from eight branches of *Eucalyptus pauciflora*.

| **Division** | **Order** | **Species** | **Size (bp)** | **LSC (bp)** | **SSC (bp)** | **IR**  **(bp)** | **HA** | **HB** | **Frequency** | **p-value** |
| --- | --- | --- | --- | --- | --- | --- | --- | --- | --- | --- |
| Angiospermae | Myrtales | *Eucalyptus pauciflora* | 159,942 | 88,787 | 18,421 | 26,367 | 1,431 | 1,459 | 0.50 | 0.62 |
|  |  | Branch tip A |  |  |  |  | 101 | 110 | 0.48 | 0.63 |
|  |  | Branch tip B |  |  |  |  | 186 | 184 | 0.50 | 0.96 |
|  |  | Branch tip C |  |  |  |  | 203 | 191 | 0.51 | 0.58 |
|  |  | Branch tip D |  |  |  |  | 132 | 114 | 0.54 | 0.28 |
|  |  | Branch tip E |  |  |  |  | 186 | 189 | 0.50 | 0.92 |
|  |  | Branch tip F |  |  |  |  | 234 | 255 | 0.48 | 0.37 |
|  |  | Branch tip G |  |  |  |  | 141 | 150 | 0.48 | 0.64 |
|  |  | Branch tip H |  |  |  |  | 248 | 266 | 0.48 | 0.45 |

**Size**: the chloroplast genome size

**HA/B**: Haplotype A/B, the number of supported reads of LSC and SSC having identical/opposite orientation.

**Frequency**: the proportion of the count of haplotype A in the count of haplotype A+B
